# Comparative analysis examining patterns of genomic differentiation across multiple episodes of population divergence in birds

**DOI:** 10.1101/202168

**Authors:** Kira E. Delmore, Juan Lugo, Benjamin M. Van Doren, Max Lundberg, Staffan Bensch, Darren E. Irwin, Miriam Liedvogel

## Abstract

Heterogeneous patterns of genomic differentiation are commonly documented between closely related populations and there is considerable interest in identifying factors that contribute to their formation. These factors could include genomic features (e.g., areas of low recombination) that promote processes like linked selection (positive or purifying selection that affects linked neutral sites) at specific genomic regions. Examinations of repeatable patterns of differentiation across population pairs can provide insight into the role of these factors. Birds are well suited for this work, as genome structure is conserved across this group. Accordingly, we re-estimated relative (*F*_*ST*_) and absolute (*d*_*XY*_) differentiation between eight sister pairs of birds that span a broad taxonomic range using a common pipeline. Across pairs, there were modest but significant correlations in window-based estimates of differentiation (up to 3% of variation explained for *F*_*ST*_ and 26% for *d*_*XY*_), supporting a role for processes at conserved genomic features in generating heterogeneous patterns of differentiation. This suggestion was reinforced by linear models identifying several genomic features (e.g., gene densities) as significant predictors of *F*_*ST*_ and *d*_*XY*_ repeatability. *F*_*ST*_ repeatability was higher among pairs that were further along the speciation continuum (i.e., more reproductively isolated), suggesting that early stages of speciation may be dominated by positive selection that is different between pairs and replaced by processes acting according to shared genomic features as speciation proceeds.

## INTRODUCTION

The integration of genomic data into research on population differentiation and speciation has led to the observation that genomic differentiation between closely related populations is often highly variable across the genome, with areas of elevated differentiation interspersed with areas of low differentiation (e.g., Nadeau et al. 2013; Renaut et al. 2013; Han et al. 2017; Vijay et al. 2017). One of the main conclusions from this observation is that speciation can proceed through a few focal changes and does not require divergence across the entire genome. This conclusion conforms to the genic view of speciation proposed by Wu (2001), but there is still considerable controversy concerning the factors that generate variation in estimates of genomic differentiation. This controversy has led to the development and extensive evaluation of two models.

The first model is termed divergence-with-gene-flow and invokes both selection and gene flow to explain heterogeneous patterns of differentiation. Specifically, this model holds that divergent selection at loci involved in population differentiation protects some regions of the genome from gene flow, elevating an otherwise homogenized landscape of differentiation (Nosil et al. 2009; Nosil and Feder 2012). The second model proposes that selection alone can generate variation in differentiation by accelerating lineage sorting at some regions of the genome. In other words, genomic regions that do not show elevated differentiation simply continue to share ancestral polymorphism (Noor and Bennett 2009; Turner and Hahn 2010; Cruickshank and Hahn 2014). We refer to this model as selection-in-allopatry and note that there are variants on this model related to when selection acts (Cruickshank and Hahn 2014; Delmore et al. 2015; Irwin et al. 2016).

One common feature of both the divergence-with-gene-flow and selection-in-allopatry models is that features of the local genomic landscape should contribute to variation in differentiation. For example, genomic regions with lower rates of recombination, higher rates of mutation and elevated gene densities can promote linked selection, defined as any form of selection that reduces variation at nearby neutral sites (Charlesworth et al. 1993; Lohmueller et al. 2011; Charlesworth 2012; Cutter and Payseur 2013; Enard et al. 2014). Linked selection can be positive, acting on new or existing mutations (genetic hitchhiking, Maynard Smith and Haigh 1974) or purifying, removing deleterious mutations from the population (background selection, Charlesworth et al. 1993). Measures of differentiation like *F*_*ST*_ include a term for within population variation and can be inflated by the reductions in variation that accompany linked selection (Charlesworth 1998). Low recombination rates make it difficult for linked neutral sites to escape the effects of new mutations via recombination. Higher mutation rates and gene densities provide more targets for selection which is also expected to be stronger in gene rich regions where mutations are more likely to have deleterious effects.

It was recently suggested that patterns of genomic differentiation will reflect features of the local genomic landscape more at later stages of speciation, as drift and background selection at these features will take time to influence differentiation (Burri 2017). Comparative analyses examining genomic differentiation across multiple population pairs are ideal for both implicating features of the local genomic landscape in generating genomic differentiation and examining their temporal effects. For example, if genomic variables are conserved across pairs, constraints imposed by processes like linked selection in these regions should generate correlated or repeated patterns of genomic differentiation. Comparative analyses are beginning to accumulate but are often limited to a closely related group of species or populations, precluding an evaluation of temporal effects and introducing statistical non-independence if a limited number of pairs are included (e.g., sticklebacks, Jones et al. 2012; sunflowers, Renaut et al. 2014; guppies, Fraser et al. 2015; songbirds, Irwin et al. 2016; copepods, Pereira et al. 2016). In addition, working at broader taxonomic scales may eliminate the role shared selective regimes play in generating repeatable patterns of differentiation, isolating the effects of genomic constraints.

Here we overcome these limitations using new and archived genomic data to estimate genomic differentiation between eight pairs of birds that span a broad taxonomic range (the most recent common ancestor to them all was ~52 MYA, http://www.timetree.org/). We look for (1) correlated patterns of genomic differentiation across these pairs (referred to as “repeatability”) and (2) an association between repeatability and the location of pairs along the speciation continuum (i.e., their level of reproductive isolation). We also (3) use linear models to implicate specific genomic features in generating repeatable patterns of genomic differentiation; these features include proxies for both recombination and mutation rates, gene density, chromosome size and proximity to chromosome ends and centromeres. We quantified the speciation continuum using hybrid zone width and genetic distance between pairs. Chromosome size, proximity to chromosome ends and centromeres may influence genomic differentiation as they have shown associations with recombination rates (Butlin 2005; Smukowski and Noor 2011). We also include linkage disequilibrium (LD) as a predictor in linear models; if linked selection is generating repeatable patterns, LD should be higher where repeatability is higher. Birds are well suited for this work as a considerable amount of information is known about speciation in this group (Price 2008) and genomic features are highly conserved across this group; birds have stable chromosome numbers, low rates of interchromosomal rearrangements, high synteny and similar recombination landscapes (Dawson et al. 2007; Griffin et al. 2007; Backström et al. 2008; Stapley et al. 2008; Ellegren 2010; Kawakami et al. 2014; Zhang et al. 2014; Kawakami et al. 2017).

Thus far we have only discussed how the local genomic landscape can affect *F*_*ST*_, a relative measure of differentiation that is inflated by reductions in within population variation. Many studies are beginning to include *d*_*XY*_ in their analyses. *d*_*XY*_ is an absolute measure of differentiation that does not include a term for within population variation. Under the divergence-with-gene-flow model of speciation, processes like linked selection at genomic features should elevate *d*_*XY*_ beyond average, background levels of absolute differentiation. Under the selection-in-allopatry model of speciation, linked selection should have no effect on *d*_*XY*_ or reduce it beyond background levels (Nachman and Payseur 2012). The latter reductions could occur in response to recurrent selection in ancestral populations that reduces variation over lineage splits and/or selective sweeps of globally adaptive alleles (e.g., Delmore et al. 2015; Irwin et al. 2016). Given increasing interest in *d*_*XY*_ and its potential to reflect the local genomic landscape, we will include it in our analyses as well. All of the population pairs included in the present study occur in the temperate region where they have likely experience periods of allopatry with glacial expansions (Hewitt 2000). Accordingly, in analyses for objective 3 where we identify specific regions that show repeatable patterns we will focus on those at the bottom of the *d*_*XY*_ distribution; *d*_*XY*_ should reflect the amount of sequence divergence that has been acquired since populations split but linked selection should reduce this parameter in some regions when gene flow is restricted between populations.

## RESULTS

The eight pairs of birds in our study include two populations of European blackcaps (*Sylvia atricapilla*), subspecies of greenish warbler (*Phylloscopus trochiloides viridanus* and *P.t. plumbeitarsus*), subspecies of willow warbler (*Phylloscopus trochilus trochilus* and *P.t. acredula*), species of stonechat (European *Saxicola rubicola* and Siberian *S. maurus*), subspecies groups of the Swainson’s thrush (coastal *Catharus ustulatus ustulatus* and inland *C. u. swainsoni*), species of flycatcher (collared *Ficedula albicollis* and pied *F. hypoleuca*), species of crow (hooded *Corvus cornix* and carion *C. corone*), and species of wood warbler (blue-*Vermivora chrysoptera* and golden-winged *V. cyanoptera* warblers; Figure 1). To gain an overview of the relationship between these pairs, we constructed a phylogeny for the group using whole-genome sequence data from all autosomal chromosomes (Figure 1). The crow is the most distantly related species; all the remaining species cluster into two groups. One group includes the greenish warbler, willow warbler and blackcap while the other includes the flycatcher, stonechat, thrush and blue/golden-winged warblers. Disregarding sister-pair relationships, the most closely related species are flycatchers and stonechats, and greenish and willow warblers. This topology is what we expected based on previous phylogenetic studies (e.g., Jetz et al. 2012, birdtree.org).

**Figure 1.**
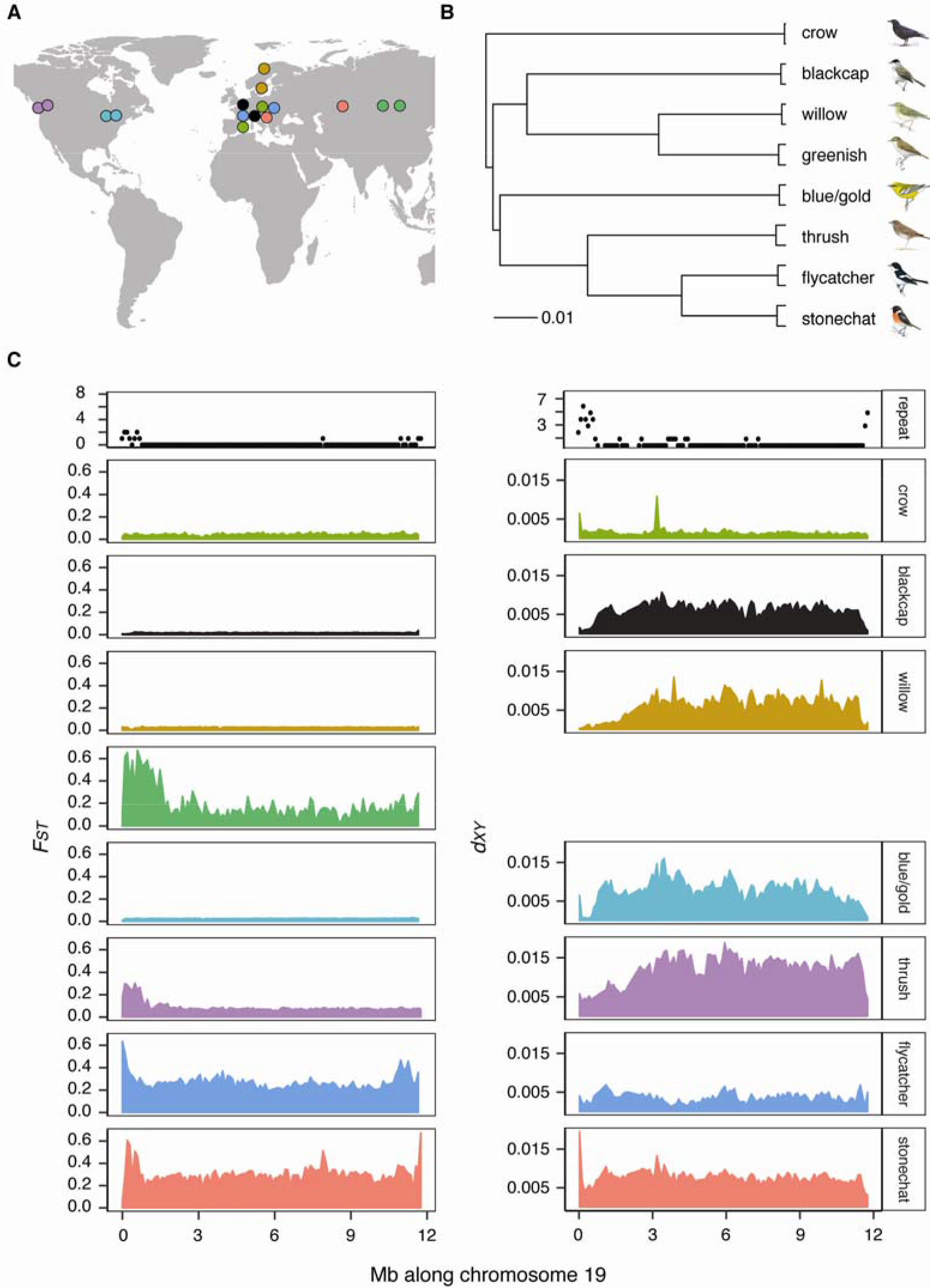
Population pairs, their (A) geographic ranges (circles as the center of sampling distributions) and (B) phylogenetic relationships (each branch has 100 bootstrapped support). Panel (C) shows windowed (100 kb) estimates of relative (*F*_*ST*_) and absolute (*d*_*XY*_) differentiation across chromosome 19 for all population pairs along with repeatability at the top, measured as the number of pairs each windows as considered an outlier in (top 5 percentile of *F*_*ST*_ distribution and bottom 5 percentile of *d*_*XY*_ distribution).

### Repeatability in patterns of genomic differentiation

To estimate repeatability in patterns of genomic differentiation across pairs we organized scaffolds from each species’ reference into chromosomes using synteny with the flycatcher and estimated *F*_*ST*_ and *d*_*XY*_ between populations in each pair using the same 100 kb windows (Figure 1). An initial comparison across pairs suggests that patterns may only be modestly consistent but stronger when considering *d*_*XY*_. We evaluated this observation by correlating windowed estimates of *F*_*ST*_ and *d*_*XY*_ across pairs. In accordance with our observation, correlation coefficients varied from −0.02 to 0.18 for *F*_*ST*_ and 0.04 to 0.51 for *d*_*XY*_ (Table 1). Squaring the highest coefficients for *F*_*ST*_ and *d*_*XY*_, these results suggest that up to 3 and 26% of the variation can be explained by correlations of *F*_*ST*_ and *d*_*XY*_ between pairs.

**Table 1.**
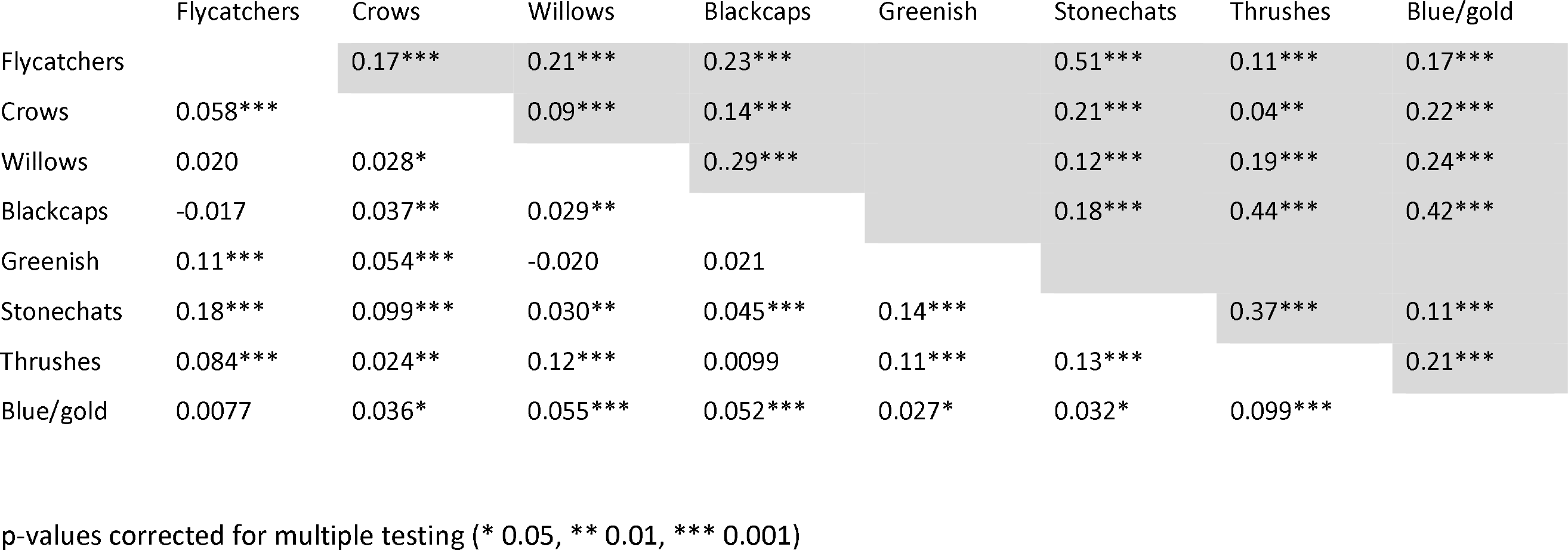
Repeatability in genomic differentiation across population pairs of birds. Values are correlation coefficients comparing windowed estimates of *F*_*ST*_ (below diagonal) and *d*_*XY*_ (above diagonal) between each set of population pairs. *d*_*XY*_ was not estimated for greenish warblers. For results based on outlier status and overlap values see Table S3.

We used a second method to quantify repeatability between pairs based on the overlap of outlier windows. We identified outlier windows for each pair as those in the top 5 percentile of the *F*_*ST*_ distribution and bottom 5 percentile of the *d*_*XY*_ distribution and compared the number of outlier windows that were shared (or overlapped) across pairs to the expected number based on the hypergeometric distribution (see Methods). Similar to results from correlations, estimates of overlap were higher and more significant for *d*_*XY*_ (Table S3a). This was also the case when we combined strings of outlier windows into peaks, acknowledging the fact outlier windows may not be independent of one another (Table S3b).

### Association between repeatability and the speciation continuum

We looked for an association between repeatability and the speciation continuum using three different measures for the speciation continuum: hybrid zone width and genetic distance based on *cytb* sequences and all autosomal sequences. Starting with hybrid zone width, we obtained estimates for each pair from the literature and assumed the more narrow a hybrid zone is the higher reproductive isolation must be (Barton and Hewitt 1985). Estimates of hybrid zone width ranged from 0 km for greenish warblers to 600 km for blue/golden-winged warblers (Table S4). We ranked pairs based on hybrid zone width and constructed a distance matrix by adding the ranks of pairs being compared (Table S4). We compared this matrix with the correlation matrices generated using windowed estimates *F*_*ST*_ and *d*_*XY*_ above. As predicted, there was a positive relationship between hybrid zone width and the correlation matrix based on *F*_*ST*_ but not *d*_*XY;*_ pairs further along the speciation continuum showed higher repeatability in *F*_*ST*_ (Mantel Test, R=0.43, p=0.03) but not *d*_*XY*_ (R=−0.20, p=0.80).

Similar to results from hybrid zone width, we documented significant positive associations between matrices based on genetic distance (*cytb* and autosomal) and repeatability in *F*_*ST*_ but not *d*_*XY*_ (Mantel Tests, *F*_*ST*_: *cytb*, R=0.36, p=0.05; autosomal, R = 0.54, p=0.02; *d*_*XY*_: *cytb*, R=−0.04, p=0.56; autosomal, R=0.12, p=0.34). Note that the associations we documented between repeatability and all three measures of the speciation continuum remain significant when controlling for phylogenetic distance (e.g., partial Mantel Tests for *F*_*ST*_ and hybrid zone width, R=0.47, p=0.02; *cytb*, R=0.36, p=0.05; autosomal, R=0.43, p=0.03) and we documented similar associations when we replaced repeatability estimated by correlation coefficients (Table 1) with values based on the overlap of outlier windows (Table S3; e.g., results for *F*_*ST*_ and hybrid zone width, R=0.29, p=0.06; *cytb*, R=0.40, p=0.01; autosomal, R=0.41, p=0.01). This is an important finding, as it suggests the associations we documented are not related to the fact that there is a greater range of *F*_*ST*_ values at later stages of speciation. Finally, note that we obtained similar results when we ran these analyses using sliding windows of 20 kb (e.g., results for *F*_*ST*_ and hybrid zone width, R=0.67, p=0.01; *cytb*, R=0.32, p=0.05; autosomal, R=0.49, p=0.03).

### Genomic features or processes as predictors of repeatability

The repeated patterns of *F*_*ST*_ and *d*_*XY*_ we documented above suggest that variation in genome-wide estimates of differentiation is influenced by conserved features of the local genomic landscape. We used generalized linear models (GLMs) to evaluate the role specific genomic features play in generating these patterns. Repeatability was quantified for each window as the number of pairs it was considered an outlier in (recall outlier status was determined using the top 5 percentile of the *F*_*ST*_ distribution and bottom 5 percentile of the *d*_*XY*_ distribution). Separate GLMs were run for each species pair using seven predictor variables (estimated for each pair separately): the proportion of GC bases (proxy for recombination rate), synonymous mutation rate (*d*_*s*_; proxy for mutation rate), gene density, LD and three variables related to where windows are located in the genome (micro- or macrochromosomes [chromosome size], proximity to chromosome ends and centromeres).

Results from these GLMs can be found in Figure 2 (with blackcap as predictor) and Table S5 (for each case as predictor). GC content, gene density, LD and proximity to both chromosome ends and centromeres were significant predictors of repeatability in *F*_*ST*_ across all species pairs. Each of these variables were positive predictors of repeatability, except GC content which was negative. In the case of position, this means that windows near the center of chromosomes showed higher repeatability. Linkage disequilibrium, proximity to centromeres and chromosome size were significant predictors of repeatability in *d*_*XY*_ across all species pairs. In the case of chromosome size, this means that windows on microchromosomes show more consistent patterns. GC content, proximity to chromosome ends, gene density and *d*_*s*_ were also significant predictors of repeatability in *d*_*XY*_ but not for all species pairs.

**Figure 2.**
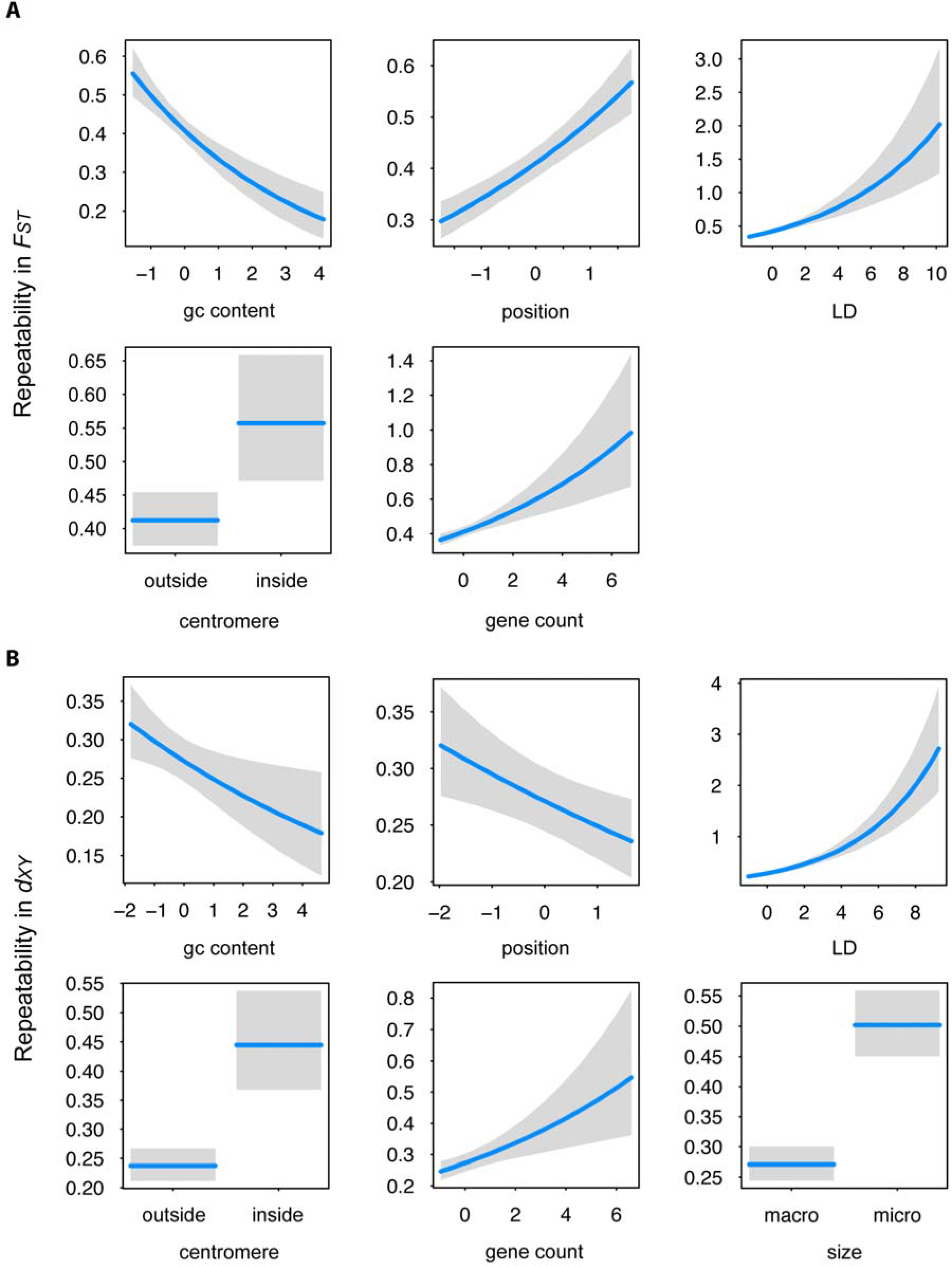
Results from GLMs examining the relationship between repeatability and predictor variables related to features of the local genomic landscape. Relationships shown are limited to significant predictor variables and results from blackcaps (results for non-significant predictor variables and the remaining population pairs and can be found in Table S5). Repeatability is estimated as the number of pairs each window was considered an outlier in (outliers are windows in the top 5 percentile of each species pairs’ distribution for *F*_*ST*_ [A] and bottom 5 percentile for *d*_*XY*_ [B]). Correlation coefficients for full models are 0.17 for *F*_*ST*_ and 0.27 for *d*_*XY*_. Parameter estimates are as follows: *F*_*ST*_, GC content −0.20 [±0.04, p<0.001], position 0.19 [±0.03, p<0.001], gene count 0.13 [±0.03, p<0.001], linkage disequilibrium 0.15 [±0.02, p<0.001], centromere 0.32 [±0.09, p<0.001]; *d*_*XY*_: GC content −0.09 [±0.04, p=0.01], position −0.08 [±0.03, p<0.001], gene count 0.11 [±0.03,p<0.001], size 0.62 (±0.08, p<0.001], linkage disequilibrium 0.24 [±0.02, p<0.001], centromere 0.63 [±0.11, p<0.001]). Each predictor is scaled and their effects are plotted with other variables held at their medians.

Note that while the predictor variables used in these models are species-specific (e.g., gene density estimate for each pair separately) the response variable is the same – a summary variable quantifying repeatability across species pairs. Accordingly, results from these models are not entirely independent.

## DISCUSSION

We studied factors influencing variation in genome-wide estimates of differentiation using genomic data from population pairs of birds that span a broad taxonomic scale. Our results support a role for (1) features of the local genomic landscape in influencing differentiation and (2) suggest that the effects of these features become more apparent later in speciation. We discuss these findings and important caveats below.

Our first objective was to determine if patterns of genomic differentiation were correlated (or repeated) across populations pairs. Our results suggest that up to 3% of the variation in *F*_*ST*_ and 26% of the variation in *d*_*XY*_ can be explained by correlations across pairs. Shared features of the genomic landscape likely contribute to these correlations. For example, linked selection (positive or purifying) is promoted by genomic features and reduces within population variation, inflating *F*_*ST*_. Many of these genomic features are conserved across birds (Dawson et al. 2007; Griffin et al. 2007; Backström et al. 2008; Stapley et al. 2008; Ellegren 2010; Kawakami et al. 2014; Zhang et al. 2014; Kawakami et al. 2017) and likely generated the repeatable patterns we observed. In support of this suggestion, recombination rates (approximated by GC content) and gene densities, two genomic features that are preserved across birds and promote linked selection were consistent predictors of repeatability in *F*_*ST*_. Genetic drift may also contribute to the correlations we documented. For example, drift in low recombination regions can cause large genomic regions to show consistently high or low *F*_*ST*_. Nevertheless, drift is generally expected to reduce genetic diversity genome-wide; it may play a localized effect if (for example) linked selection reduces *N*_*e*_ in areas of low recombination but this scenario is not independent of selection (reviewed in Charlesworth 2009). Note that the amount of variation explained by correlations across pairs is not as high as other studies (e.g., 49-77% of the variation in *F*_*ST*_ explained by correlations across subspecies pairs of greenish warblers in Irwin et al. 2016). Nevertheless, most of these studies focus on pairs that are much more closely related than those in the present study, such that pairs share more genomic features and are subject to similar selective forces.

Our second objective was to determine if there was a positive association between repeatability and the location of pairs along the speciation continuum. We measured the speciation continuum using hybrid zone width and genetic distance and found that pairs with narrower hybrid zones and greater genetic distances exhibited more similar patterns of *F*_*ST*_. The latent effects of drift and background selection may explain this pattern. Specifically, when population divergence begins, allele frequencies will be roughly equal and *F*_*ST*_ will be close to zero. Drift and selection will start acting on standing genetic variation and any new mutations that arise. The effects of drift and background selection may take time to accumulate. Accordingly, positive selection may be the primary force affecting differentiation early in speciation (Burri 2017). If the selective context of speciation is different for the pairs under study, this should lead to less repeatable (or correlated) patterns of differentiation. As speciation proceeds, the effects of drift and background selection will become more pronounced and result in the landscape of differentiation reflecting genomic features that promote these processes (along with positive selection and genetic hitchhiking) more directly (Burri 2017). Note that the beginning stages of speciation may be less repeatable even without different selective pressures. For example, there may be more than one way to respond to selection and the chance positive selection affects the same genomic region may be low. In addition, beneficial mutations are rare (Eyre-Walker and Keightley 2007); during the time it takes for positive selection to act, the regions where differentiation becomes elevated in response to drift and background selection may be unpredictable (Burri 2017).

Thus far we have focused mainly on results for *F*_*ST*_; results for *d*_*XY*_ require careful explanation. Concerning the repeatable patterns we documented (first objective, up to 26% of the variation in *d*_*XY*_ explained by correlations across pairs), at the outset we argued that speciation in the pairs we studied would have been punctuated with periods of allopatry as all pairs occur in the temperate region where glacial advances would have isolated populations in different refugia (Hewitt 2000). Under this scenario (i.e., without gene flow), *d*_*XY*_ should reflect the amount of sequence divergence that has been acquired since populations split and linked selection should have no effect on *d*_*XY*_ or reduce it (e.g., if recurrent selection in ancestral populations removes variation from populations prior to their split; Nachman and Payseur 2012; Cruickshank and Hahn 2014). In support of this finding, several genomic features including gene densities and chromosome size predicted repeatable patterns at the bottom of the *d*_*XY*_ distribution. Nevertheless, it is important to note that linked selection can also elevate *d*_*XY*_, especially during periods of secondary contact where gene flow would have occurred and thus some of the repeatable patterns we documented may also be related to elevated *d*_*XY*_ (Nachman and Payseur 2012).

Continuing with *d*_*XY*_ repeatability (i.e., results for the first objective), correlation coefficients were higher for *d*_*XY*_ than *F*_*ST*_, with the highest correlation coefficient for *d*_*XY*_ being more than twice that for *F*_*ST*_ (0.18 vs. 0.51). This finding could be related to the fact that *d*_*XY*_ reflects processes that have accumulated over multiple speciation events (see below for additional explanation) while *F*_*ST*_ mainly reflects process in extant populations. Accordingly, if population pairs are sampled too early in speciation, *F*_*ST*_ may not reflect local genomic features yet (Burri 2017). This suggestion follows from the argument described above about the latent effects of drift and background selection. Nevertheless, additional explanations for increased *d*_*XY*_ repeatability are also possible. For example, *d*_*XY*_ shows a strong relationship with mutation rates (Geneva et al. 2015; Rosenzweig et al. 2016). Accordingly, much of the pattern we documented may be related solely to variation in mutation rates. It is also important to note that *d*_*XY*_ is estimated using far more sites than *F*_*ST*_ (variant and invarant vs. just variant for *F*_*ST*_). Accordingly, these estimates may be more precise, leading to stronger correlation coefficients.

Finally, while we found a correlation between the speciation continuum and repeatability in *F*_*ST*_ (second objective) we did not document this association for *d*_*XY*_. Again, this finding may be related to the fact that *d*_*XY*_ reflects processes that have accumulated over multiple speciation events while *F*_*ST*_ mainly reflects process in extant populations, including speciation. For example, if the recombination landscape has remained the same for millions of years, recurrent linked selection in areas of low recombination has likely been reducing variation over the same time period and these reductions will be passed down over speciation events (Burri 2017). Under this scenario, it will not matter what stage of differentiation the population pairs under study are at, these reductions will be reflected in estimates of *d*_*XY*_. As we have already discussed, there are situations where *d*_*XY*_ will reflect processes in extant populations (especially if gene flow is occurring) but the underlying effect of ancestral diversity appears to override any effect these processes have on *d*_*XY*_ repeatability and the speciation continuum.

We used a comparative analysis to provide insight into how heterogeneity in both relative and absolute differentiation are generated and change during the process of speciation, implicating processes at conserved genomic features in generating variation in both measures of differentiation and providing evidence that these processes have a greater effect on relative estimates of genomic differentiation later in speciation. By working at such a broad taxonomic scale we were able to reduce the effect shared selective regimes may play in generating repeated patterns of differentiation. By including population pairs with good information on reproductive isolation, we were able to provide some of the first empirical evidence for the latent effects of drift and background selection on relative differentiation and highlight the importance of the selective context (i.e., positive selection and genetic hitchhiking) early in speciation. Our results support the general sentiment (e.g., Ravinet et al. 2016) that caution must be used when interpreting patterns of differentiation, as processes other than positive selection can influence differentiation. In addition, there is increasing interest in applying additional measures of differentiation beyond *F*_*ST*_ to speciation genomics but as highlighted by the caveats we discuss for *d*_*XY*_, careful consideration must be used when applying them to this topic.

## METHODS

### Study Species and Datasets

We searched the Sequence Read Archive (https://www.ncbi.nlm.nih.gov/sra) and European Nucleotide Archive (http://www.ebi.ac.uk/ena) for genomic data collected from birds. We limited our search to species for which a draft reference genome had been assembled for the target species or one that was closely related. This search resulted in the inclusion of eight pairs (Table S1). The only pair we did not have a reference genome for was the *Vermivora* warblers but a reference for the closely related yellow-rumped warblers (*Setophaga coronata*) is available and was used in the original publication for these data (Toews et al. 2016). All pairs are from the order Passeriformes (perching birds or songbirds) and breed in temperate regions (Figure 1).

### Generating consensus reference genomes

The reference genomes we acquired were all assembled into scaffolds, except the collared flycatcher’s genome, which was organized into chromosomes based on linkage map for the species and synteny with zebra finch (Ellegren et al. 2012). Accordingly, we used this genome to ensure all windows compared across species were orthologous. To maintain chromosomal synteny, we aligned the scaffolds of each genome individually against the flycatcher genome with SatsumaSynteny (default parameters; Grabherr et al. 2010). We then used bash scripts to parse the output, obtaining information on the order and orientation of query scaffolds and conducted a final alignment with the LASTZ plugin in Geneious (Harris 2007; Kearse et al. 2012). We merged these scaffolds into pseudochromosomes by calling the query base where alignments occurred and Ns in the presence of a gap. Details on consensus genome coverage can be found in Table S2.

### Constructing phylogenetic network

We used ANGSD (Korneliussen et al. 2014) to obtain consensus fasta sequences for populations from each species pair (-doFasta 2-doCounts 1-minQ 20-setMinDepth 10) and IQ-TREE (Nguyen et al. 2016) to generate a distance matrix based on these sequences. We used the resulting distance matrix to construct a phylogenetic tree using the UPGMA clustering method in PHYLUCE (neighbor, http://evolution.genetics.washington.edu/phylip/doc/neighbor.html).

### Estimating differentiation

We focused on SNPs for the present study and used a common reference-based bioinformatics pipeline to call them. Details can be found in Delmore et al. (2015) and Delmore et al. (2016). Briefly, we trimmed reads with trimmomatic (TRAILING:3 SLIDINGWIND0W:4:10 MINLEN:30) and aligned them to consensus genomes using bwa *mem* (Li and Durbin 2009) using default settings. We then used GATK (McKenna et al. 2010) and picardtools (http://broadinstitute.github.io/picard) to identify and realign reads around indels (*RealignerTargetCreator*, *IndelRealigner*, default settings) and removed duplicates (*MarkDuplicates*, default settings) for all datasets except greenish warbler which consisted of GBS data.

We used two estimates of differentiation in our study: *F*_*ST*_ and *d*_*XY*_. We estimated *F*_*ST*_ for datasets comprised of individuals using ANGSD, estimating site frequency spectrums for each population separately (-dosaf 1, -gl 1, -remove_bads, -unique_only, -minMapQ 20, -minQ 20, -only_proper_pairs 1, -trim 0) and using these to obtain joint frequencies spectrums for population pairs. These joint frequency spectrums were then used as priors for allele frequencies at each site to estimate *F*_*ST*_. For datasets comprised of pools we estimated *F*_*ST*_ using Popoolation2 (Kofler et al. 2011; -min-coverage 30 for Swainson’s thrushes and 10 for stonechats, -min-count 3, -minq 20). We summarized *F*_*ST*_ into windows of 100 kb and limited analyses to windows with data from all pairs. We excluded the Z chromosome from all analyses as some of the pairs included females where systematic biases related to coverage could affect estimates of differentiation.

We estimated *d*_*XY*_ for datasets comprised of individuals using ANGSD as well. First, we estimated allele frequencies at each SNP for both populations of each pair combined - doMajorMinor 4, -doMaf 2, -gl 1, -doCounts 1, -remove_bads, -unique_only, -minMapQ 20, -minQ 20, -only_proper_pairs 1, -trim 0, -SNP_pval 1e-6). We then reran the program by population using only the SNPs that passed the previous step, to ensure SNPs fixed in one population were not lost. Once we had these allele frequencies, we estimate *d*_*XY*_ at each SNP using a script provided with ANGSD (https://github.com/mfumagalli/ngsPopGen//scripts/calcDxy.R) and as (p1*(1−p2))+(p2*(1−p1)) where p is the allele frequency of a given allele in populations 1 and 2, respectively and averaged these values in the same 100 kb windows used for *F*_*ST*_. Estimates of *d*_*XY*_ by SNP have to be normalized by the number of sites (variant and invariant) in a window. We obtained these values using ANGSD to estimate read depth at all sites (-doCounts 1, -dumpCounts 1, -remove_bads, -unique_only, -minMapQ 20, -minQ 20, -only_proper_pairs 1, -trim 0) and excluded sites with coverage less than three times the sample size and more than three times the average coverage to ensure roughly three reads per individual and exclude sites that may have mapping problems resulting from copy number variants. Analyses were limited to windows with data from all pairs and windows with more than 5000 callable sites, as *d*_*XY*_ can be highly variable with small sample sizes and coverage (e.g., Clarkson et al. 2014). This filter precluded the use of greenish warblers in analyses of *d*_*XY*_ as data for this pair were derived from reduce-representation sequencing and did not have high coverage in windows of 100 kb. This was not a problem for the original publication (Irwin et al. 2016) as windows were defined by SNPs rather than base pairs.

For stonechats and thrushes, for which we used pooled sequencing data, we calculated *d*_*XY*_ with a custom script. We excluded sites with coverage below 10 for stonechats and below 30 for thrushes. We estimated *d*_*XY*_ by multiplying allele frequencies for each base as above and averaging across sufficiently covered bases in each window.

### Estimating repeatability in patterns of genomic differentiation

We estimated overall repeatability between pairs by correlating windowed-estimates of differentiation across pairs. We also estimated repeatability using information on outlier status and overlap. Specifically, we identified outlier windows for each species pairs as those above the 5% quantile for *F*_*ST*_ and below the 5% quantile for *d*_*XY*_. For each comparison across pairs (e.g., flycatcher to stonechat, flycatcher to greenish warbler, etc.), we counted the number of outlier windows that were shared (or overlapped) and compared this to the expected number of overlapping windows using the hypergeometric distribution, assuming that each window had an equal probability of being considered an outlier. We used these values (observed and expected) to calculate z scores for each comparison and calculated one-sided p-values (i.e., the probability of obtaining an overlap value as extreme or more extreme than our observed value). Z scores are effect sizes that correspond to the number of standard deviations above the average expectation in each comparison, allowing for direct comparison across studies. Note that it is also possible that the outlier windows we identified are not independent of one another. Accordingly, we modified our overlap approach by combining outlier windows into peaks if they occurred next to one another and using the number of peaks as our estimate of overlap. We used a permutation test to quantify the significance of these values, holding the number and size of outlier regions constant while randomly permuting their location 1000 times and calculating one-sided p-values again (Van Doren et al. 2017).

### Placing pairs along the speciation continuum

We used three methods to place pairs along the speciation continuum (i.e., to quantify the level of reproductive isolation), starting with the width of hybrid zones, which we obtained from the literature. The more narrow a zone the higher the reproductive isolation (Barton and Hewitt 1985; Moore and Dolbeer 1989; Paradis et al. 1998). If hybridization is extremely rare (e.g., with the greenish warblers), we set the width as zero. Note the use of hybrid zone width as a measure for reproductive isolation assumes all hybrid zones are at an equilibrium between selection and dispersal (Barton & Hewitt 1985) and that dispersal distances are similar across pairs. We also used genetic distance within pairs, which we obtained by aligning *ctyb* sequences (downloaded from NCBI https://www.ncbi.nlm.nih.gov/) using MEGA as *p*-distance (the proportion of nucleotide sites that differ between groups) and the distance matrix generated by IQTREE for autosomal chromosome alignments (see “Consructing phylogenetic network”). We used Mantel tests to compare distance matrices quantifying the speciation continuum with distance matrices quantifying repeatability. We accounted for small sample sizes in these tests by using permutation tests to quantify significance and the non-parametric Spearman rank correlation coefficient.

### Measuring genomic features and processes and their effect on repeatability

We looked at the relationship between repeatability and seven structural features of the genome: recombination rate, mutation rate, gene density, chromosome size, proximity to chromosome ends and centromeres and linkage disequilibrium. We estimated these features for each species and ran separate generalized linear models with repeatability as the response variable with a Poisson distribution for each species.

We used GC content as a proxy for recombination. Recombination affects the patterns of local base composition via the unbalanced transmission of ‘strong’ (GC) over ‘weak’ (AT) alleles at double-strand breaks (Mugal et al. 2015). This process is termed GC-biased gene conversion and direct support was recently presented in birds (Smeds et al. 2016). Positive correlations between recombination and GC content have also been documented in birds (Kawakami et al. 2014; Burri et al. 2015). Synonymous mutations occur in the exon of genes but have no effect on the sequence of amino acids. The use of *d*_*s*_ for mutation rate analysis assumes these sites do not experience selection (Eyre-Walker and Keightley 1999). We used a phylogenetic framework to obtain these estimates; details can be found in the Supplementary Methods. Briefly, we annotated each consensus genome with MAKER and identified potential homologues for high quality transcripts (AED < 0.05) using a Blastn search against all transcripts from the flycatcher (flycatcher was searched against zebra finch). We aligned codons from each pair of sequences using PRANK (http://wasabiapp.org/software/prank) and calculated *d*_*N*_/*d*_*s*_ with PAML v4.8. All *d*_*N*_/*d*_*s*_ calculations were performed pairwise, comparing all the species with the flycatcher and this in turn, compared to zebra finch. Estimates of *d*_*s*_ were extracted for GLMs. We used PLINK 1.9 (Chang et al. 2015) to estimate linkage disequilibrium for one population from each pair (the same population for which we had a reference genome), as the squared correlation coefficient (*r*^2^) between pairs of SNPs. SNPs were output from ANGSD using the same filters described above for *F*_*ST*_ and *d*_*XY*_ and including an additional filter for minor allele frequency, requiring SNPs have minor allele frequencies greater than 0.05. PLINK was run with the command line with the command line ‘--ld-window 100 --ld-window-kb 100 --ld-window-r2 0’ to limit the analyses to SNPs with fewer than 100 variants between them and no more than 100 kilobases apart and report pairs with *r*^2^ values below 0.2 as well. We determined the midpoints for all SNP pairs, binned them into the same 100 kb windows used for *F*_*ST*_ and *d*_*XY*_ and calculated average values for each window.

Avian genomes are composed of micro- and macrochromosomes. We considered microchromosomes those that are less than 20 Mb (Ellegren 2013) and macrochromosomes those that are greater than 40 Mb. We identified the position of each window along the chromosome by dividing chromosomes in half and generating a variable that increased by 1 from each end to the middle of a chromosome. We standardized this measure by dividing these values by half the number of windows on each chromosome. We inferred the location of centromeres using methods employed by Ellegren et al. (2012) and Delmore et al. (2015). Specifically, we identified FISH probes from Warren *et al.* (2010) on either side of centromeres in the zebra finch genome and used NCBI’s blastn (Altschul et al. 1990) to find their location in our genome. We considered sequences between FISH probes as “centromeric regions” and note this method only gives us a rough approximation for the location of centromeres.

## ACKNOWLEDGEMENTS

We acknowledge funding from the Max Planck Society and NSERC and discussions with Diethard Tautz, Diana Rennison, Kieran Samuk, Greg Owens, Sara Miller, Barbara Helm, Matthias Weissensteiner and Niclas Backström.

## AUTHOR CONTRIBUTIONS

KD and MLi conceived and designed the study, all authors contributed data. KD conducted all analyses excluding the estimation of *d*_*XY*_ for pooled datasets and repeatability using peaked overlap values (BV). JL helped with the construction of consensus genomes and estimates of *d*_*s*_ and MLu with the estimation of *d*_*XY*_ for individual datasets. KD wrote the manuscript with comments from all authors.

## DATA ACCESSIBILITY

Accession numbers for all genomic data used in this paper can be found in Table S1.

